# A framework for the analysis of symmetric and asymmetric divisions in developmental processes

**DOI:** 10.1101/010835

**Authors:** Pau Rué, Yung Hae Kim, Hjalte List Larsen, Anne Grapin-Botton, Alfonso Martinez Arias

## Abstract

Animal tissue development relies on precise generation and deployment of specific cell types into tissue sub-structures. Understanding how this process is regulated remains a major challenge of biology. In many tissues, development progresses through a sequence of dividing progenitor cells, each with decreasing potency, that balance their growth and differentiation. Dividing progenitor cells thus face a decision on whether their offspring shall differentiate or self-renew. This results in three possible modes of division (symmetric self-renewing, symmetric differentiating, and asymmetric) all of which have been observed in developing animal tissues. In some instances, the frequencies of occurrence of these division modes are incompatible with the possibility that sibling cells take the decision to differentiate independently of each other. Rather, an excess of symmetric divisions, both proliferating and differentiating, is usually observed in so far no general mechanism by which this fate entanglement takes place has been put forward. Here we propose a simple model of progenitor priming that provides a rationale on how the fate of sibling cells might be linked. Analysis of the model suggests that commitment to the cycle completion of cells primed for differentiation might be the cause of the observed excess of symmetric divisions. The model presented is applicable to a broad range of developmental systems and provides a testing framework to explain the dynamics of cell division and differentiation are related.

## Introduction

The development of an organism relies on the expansion of one cell through multiple rounds of cell division and the controlled differentiation of these cells over the time. Tissues and organs have specific sizes and for these reason, there has to be a balance between the expansion of a cell pool and its differentiation. A common strategy to achieve this balance is to set a population of multipotent progenitors for a particular tissue or organ which first multiply to a critical size from which they can differentiate into the cell types of the system. In many instances there is a shift over time from progenitor proliferative divisions to differentiating divisions. The development of the Central Nervous System (CNS) is a well characterized example of this behaviour/strategy (Caviness, Takahashi, and R. Nowakowski 1995; Hardwick and Philpott 2014; McConnell 1995; Rakic 1995; Rapaport et al. 2004; Takahashi, R. S. Nowakowski, and Caviness 1996). The early embryonic development segregates a structure, the neural plate, made up of progenitors which begin to proliferate to amplify. Over time, the divisions begin to produce differentiating cells which exit the cell cycle and accumulate developing their specific characteristics. A common way of representing these processes is to consider that the division of a Progenitor (*P* ) can give rise to two equal cells, which can be either *P* or Differentiating (*D*) cells (Livesey and Cepko 2001; Takahashi, R. S. Nowakowski, and Caviness 1996) or two different cells, *P* and *D*. The first kind of division is called symmetric and the second one asymmetric. We shall use the term asymmetric in the sense of a division that generates two different cells, independently of whether this is due to distinct cues introduced during mitosis or afterwards. Asymmetric cell divisions maintain progenitor cell numbers constant (each dividing cell is replaced by one daughter), whilst symmetric divisions alter the number of cells in the proliferative pool favouring or not the differentiating pool. In the case of the CNS what is known is that during development an initial prevalence of *P → P P* divisions is shifted to *P → DD* after going through phases with defined proportions of *P P*, *P D* and *DD* divisions (Caviness, Takahashi, and R. Nowakowski 1995; Götz and Huttner 2005; Livesey and Cepko 2001; Takahashi, R. S. Nowakowski, and Caviness 1996).

Advances in genetic labelling and lineage-tracing techniques have made it possible to quantitatively measure the proliferation and differentiation events at the single cell resolution (Blanpain and Simons 2013; Kretzschmar and Watt 2012). This progress has been accompanied by the development of theoretical models of stochastic cell division (Klein and Simons 2011). These models have proved useful in analysing homeostatic systems and are slowly being adapted for use in developmental (He et al. 2012; Lescroart et al. 2014) and regeneration (Centanin et al. 2014; Hara et al. 2014) systems. Despite their utility, these focus on the distributions of clone sizes (Klein and Simons 2011) but do not address the question of how is the distribution of division modes shaped.

While a great deal is known about the mechanisms that promote asymmetric cell division (e.g., asymmetric segregation of determinants linked to spindle orientation, uneven inheritance of cytoplasmic content or activation of different genetic programmes under the control of the environment (Gönczy 2008; Knoblich 2008)), systems with enriched amounts of symmetric divisions still ask for mechanistic ellucidations.

Here, we develop a simple model of cell differentiation that provides a framework to study the evolution of developing tissues in terms of cell division and differentiation. The model relies only on two parameters that can be inferred from the experimental data, can account for an excess in symmetric divisions in developing systems. Furthermore, it lays out several testable predictions and provides quantitative intuition on the role of each division mode on the growth dynamics. We provide a few experimental examples where the model predictions are validated and suggest that this mechanism of cell division and differentiation might be general in developing tissues.

## Results

### Cell proliferation and cell differentiation: a framework

When a cell, *P*, divides it can give rise to two types of cells: either a copy of itself (*P* ) or a different one (*D*), This translates into three types of divisions that are commonly known as symmetric *P P* or *DD*) and asymmetric (*P D*) (Fig. 1A). In systems in which one can track the individual cells, it is possible to account of each and every type of division. However, in large populations with asynchronous divisions that can only be accounted for by cell labelling methods such the clonal pair-cell analysis (Li et al. 2003), this classification provides a framework to interpret the dynamics of growth and differentiation. Indeed, the experimental measure of the relative proportions of each of the three modes of division *P*_*PP*_, *P*_*PD*_ and *P*_*DD*_ provides a straightforward way to estimate the probability of cell differentiation (*q*). This probability indicates the so called *Q* fraction (Caviness, Takahashi, and R. Nowakowski 1995; Livesey and Cepko 2001; Takahashi, R. S. Nowakowski, and Caviness 1996), i.e., the fraction of newborn cells that will eventually leave the progenitor pool, *P*, and differentiate and can thus be calculated as the contribution of all cells stemming from *DD* divisions plus half of the cells from *P D* divisions:

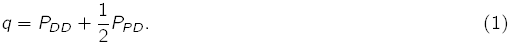

**Figure 1.**
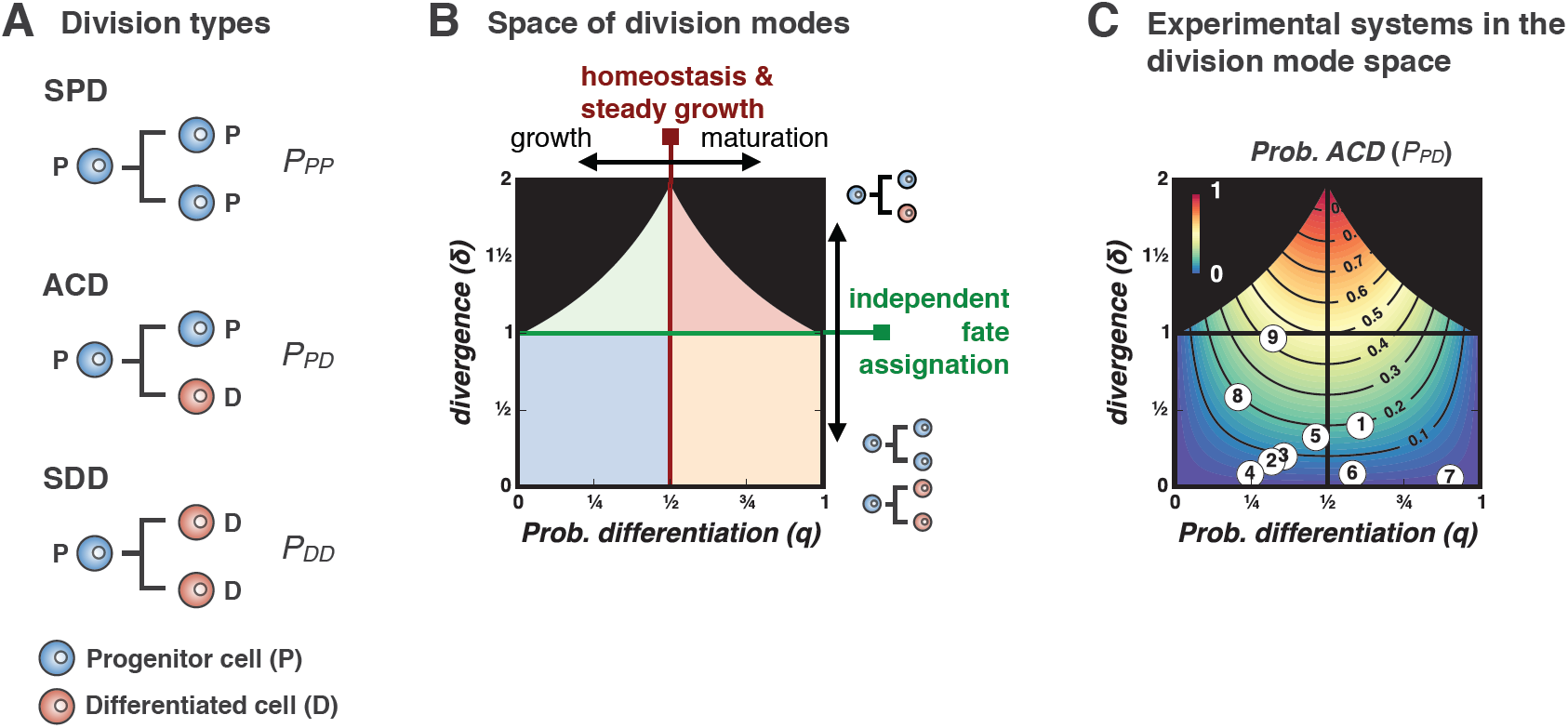
Proliferation and differentiation: cell division modes and the q − δ phase space. **A)** Proliferative (*P* ) cells can divide in one of three ways: symmetrically generating two proliferative cells (*P P* ), asymmetrically, giving rise to one proliferative and one differentiated cell (*P D*) or symmetrically into two differentiated cells (*DD*). **B)** The proportions of each division mode (*P*_*PP*_, *P*_*PD*_ and *P*_*DD*_) establish a dynamic regime of cellular proliferation and differentiation that is characterized by the fraction *q* of differentiated cells, and the *divergence*, *δ*. The first parameter (*q*) is the probability of differentiation and the second (*δ*) measures whether sibling cells tend to differentiate in conjunction through symmetric divisions (*δ <* 1), they adopt independent fates by means of asymmetric divisions (*δ >* 1) or they commit to differentiation independently of each other (*δ* = 1), **C)** Developmental and experimental systems can be described by these two parameters, and can be mapped onto the *q − δ* space as points (if they keep *q* and *δ* constant) or as trajectories (if these parameters change over time). Numbers displayed in the phase space correspond to the experimental systems summarized in Table 1.

**Table 1.**
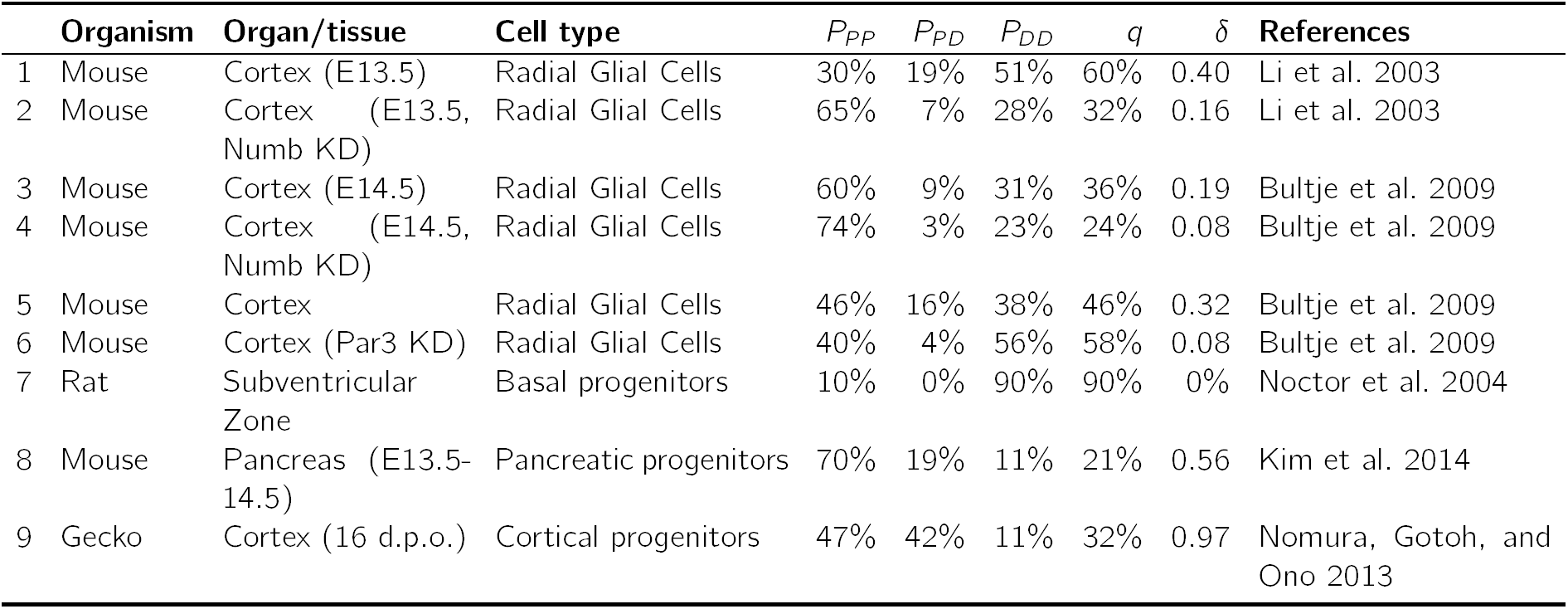
Representative developmental systems for which the frequencies of each division mode (*P P*, *P D* and *DD*) have been obtained experimentally. These are represented in division mode phase space in Fig. 1C.

| | Organism | Organ/tissue | Cell type | $P_{PP}$ | $P_{PD}$ | $P_{DD}$ | $q$ | $\delta$ | References |
| --- | --- | --- | --- | --- | --- | --- | --- | --- | --- |
| 1 | Mouse | Cortex (E13.5) | Radial Glial Cells | 30% | 19% | 51% | 60% | 0.40 | Li et al. 2003 |
| 2 | Mouse | Cortex (E13.5, Numb KD) | Radial Glial Cells | 65% | 7% | 28% | 32% | 0.16 | Li et al. 2003 |
| 3 | Mouse | Cortex (E14.5) | Radial Glial Cells | 60% | 9% | 31% | 36% | 0.19 | Bultje et al. 2009 |
| 4 | Mouse | Cortex (E14.5, Numb KD) | Radial Glial Cells | 74% | 3% | 23% | 24% | 0.08 | Bultje et al. 2009 |
| 5 | Mouse | Cortex | Radial Glial Cells | 46% | 16% | 38% | 46% | 0.32 | Bultje et al. 2009 |
| 6 | Mouse | Cortex (Par3 KD) | Radial Glial Cells | 40% | 4% | 56% | 58% | 0.08 | Bultje et al. 2009 |
| 7 | Rat | Subventricular Zone | Basal progenitors | 10% | 0% | 90% | 90% | 0% | Noctor et al. 2004 |
| 8 | Mouse | Pancreas (E13.5-14.5) | Pancreatic progenitors | 70% | 19% | 11% | 21% | 0.56 | Kim et al. 2014 |
| 9 | Gecko | Cortex (16 d.p.o.) | Cortical progenitors | 47% | 42% | 11% | 32% | 0.97 | Nomura, Gotoh, and Ono 2013 |

Its inverse, *q*^−1^ can be regarded as a measure of the average number of divisions required in order to obtain a differentiated cell. Hence, if *q* is less than 0.5, the portion of new cells that will differentiate (*D*) will be less than that of cells that will continue proliferating and in this case the progenitor pool (*P* ) will expand thus establishing a regime of exponential growth. For *q* exactly 0.5 the system attains a balance or homeostasis of the progenitor pool, as both proliferation and differentiation will equilibrate. In this case and in the absence of noticeable cell loss, the differentiated pool will increase steadily at a linear pace. Conversely, in cases in which loss of cells cannot be neglected (as in most homeostatic systems), both the proliferative and differentiated cell populations level off and are maintained in equilibrium. Finally, for *q* greater than 0.5, on average, the number of progenitors diminishes after each division (i.e., the average number of progenitors generated per dividing progenitor, 2(1 *− q*), is less than one). Therefore, regardless of the particular links existing between cell division and differentiation, the proportion of *D* cells in the population (i.e., the *Q* fraction), as measured from the division modes, is a direct indicator of whether a system is expanding, balanced or maturing.

It is important to note, however, that the relation between the *Q* fraction and the frequency of each division type is not unambiguous, i.e., there are infinitely many combinations of *P*_*PP*_, *P*_*PD*_ and *P*_*DD*_ that lead to the same probability *q* and thus given only the probability of differentiation one can not infer the frequencies of each division type. Indeed, *q* can be used to calculate the linkage between cell division and cell differentiation. If cell fate and cell division were independent events sibling cells would adopt fates independently of each other and the proportions expected for each division mode in a population would follow the binomial expansion. This situation is reminiscent of that of alleles following the Hardy-Weinberg equilibrium in a population (Hardy 1908):

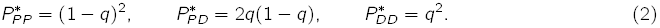

As with Hardy-Weinberg principle, lack of equilibrium implies a linkage between differentiation and division, and this can be evaluated statistically by means of Fisher’s exact test or Pearson’s *χ*^2^-test(Fisher 1922), provided the absolute numbers of each division mode are recorded.

Recent advances in genetic labelling of allowed the observation and measurement of individual cell division events (Blanpain and Simons 2013; Kretzschmar and Watt 2012). Contrary to what had been suggested in the early days (Livesey and Cepko 2001; Takahashi, R. S. Nowakowski, and Caviness 1996), these results in general indicate that the *P P/P D/DD* proportions diverge from equilibrium (see Table 1). Such deviation from the equilibrium proportions can be measured by just one factor. Take, for instance, the real fraction of *P D* divisions, *P*_*PD*_, which might differ by a Δ = *P*_*PD*_ −2*q*(1 *− q*), where the fraction *q* can be estimated experimentally using equation (1). The deviation in the other two fractions will then be 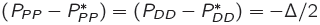. A more convenient measure, as we will see below, is the relative deviation, measured as a fold change:

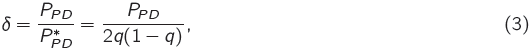

Note that *δ* is unambiguously related to Δ (*δ* = 2*q*(1 *− q*)(*δ −* 1)); it is only defined for 0 *< q <* 1 and for a given *q* can only take values in the range 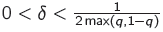because the fraction of *P D* division cannot exceed 2*q* nor 2(1 *−q*). That is to say, the requirement for a fast growth (and low *q*) or fast maturation (and high *q*) impose a limit in the maximum number of asymmetric divisions the system can sustain, as these do neither contribute to the exponential expansion nor to the fast differentiation.

For divergence *δ >* 1, the number of observed asymmetric divisions exceeds the expected and this might be suggestive of a particular mechanism of asymmetry promotion (e.g., uneven splitting of cytosolic content, planar cell polarity, epithelial polarity and loss of contact with the basal lamina, etc.). On the other hand, *δ <* 1 might hint at the existence of mechanisms preserving identical fates in sibling cells. The *divergence* parameter *δ* is therefore complementary to the *q* fraction and both define the phase space of division modes (see Fig. 1B).

The division mode space, which consists of the axes *q* and *δ* and encompasses all possible combinations of *P*_*PP*_, *P*_*PD*_ and *P*_*DD*_ values in the region described above (Fig. 1B). The line *q* = 1*/*2 separates between expanding and maturing systems, and the line *δ* = 1 does so with systems rich in asymmetric vs. symmetric divisions. The phase space thus allows to represent any stage of a developing or homeostatic system provided it can be defined by the three division modes where their proportions are maintained over a period of time.

Table 1 and Figure 1C illustrate a few examples that have been experimentally characterized in terms of *P P*, *P D* and *DD* divisions. For instance, during cortex development most cells divide symmetrically to expand the pool, and only a few undergo asymmetric division that seem to be regulated by Notch signalling, Numb/Numb-like, and the mammalian cell polarity determinant mPar3 (Bultje et al. 2009; Li et al. 2003). Disruption of either Numb or mPar3 effectively suppresses the asymmetric divisions (*δ ≈* 0, Fig. 1C). These changes in the division mode are captured by the parameter *δ* (Fig. 1C). Another example is found in mouse pancreas development. We have recently shown that during the expansion phase by bipotent pancreatic progenitors (E13.5-14.5), these show a division pattern biased towards symmetric divisions (*δ* = 0.56, Fig. 1C) (Kim et al. 2014).

### Dynamics of cell proliferation and differentiation in developing tissues: a model of stochastic cell priming

An intriguing feature of the division mode space depicted in 1C is that most expanding systems are biased towards high proportions of symmetric divisions.

Hereafter we focus on such developing systems and propose a model which explains this overabundance of symmetric divisions. The model (Fig. 2A), which accounts for cell proliferation and differentiation, considers three distinct cellular types, namely

**Figure 2.**
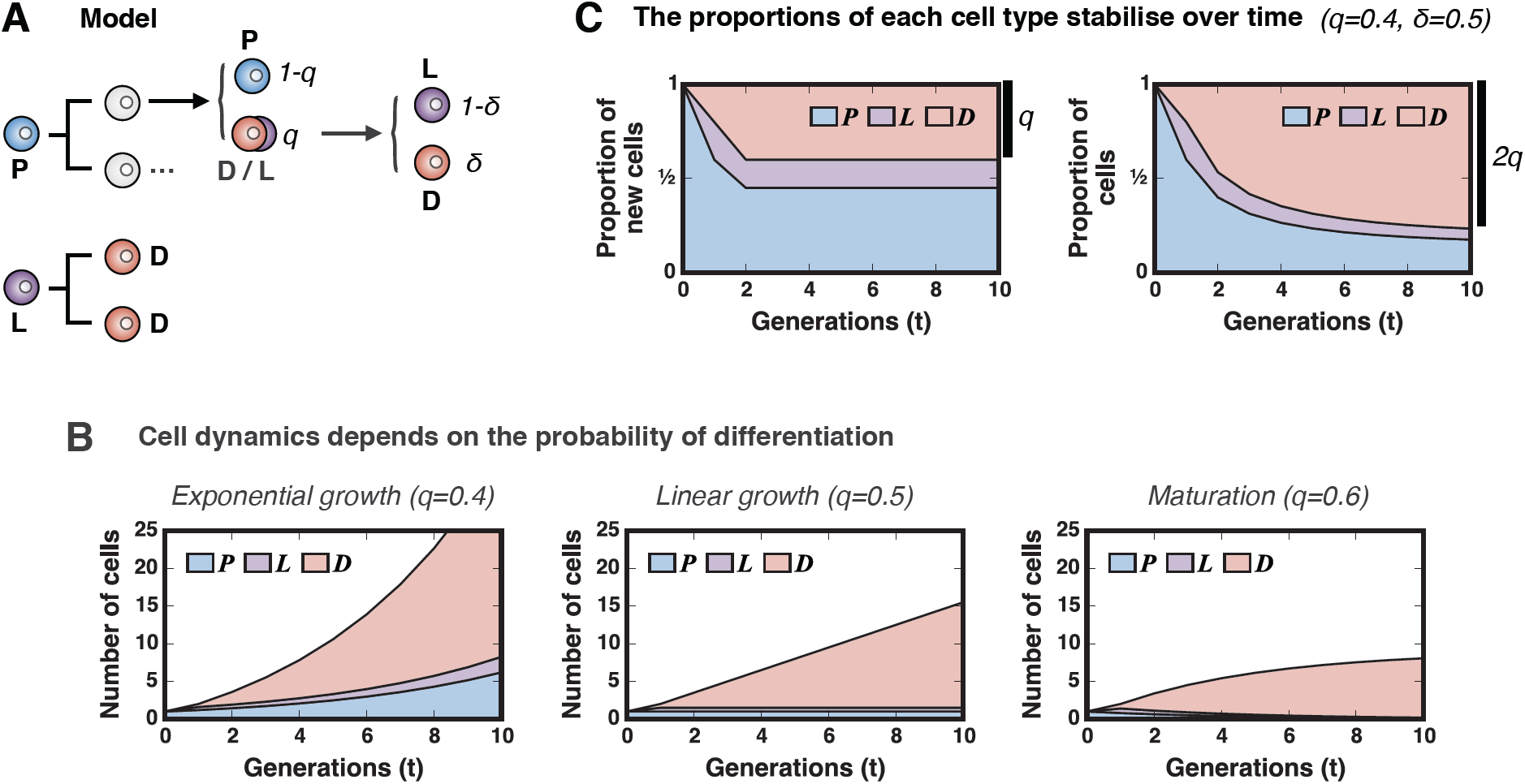
Model of cell proliferation and differentiation. **A)** Schematic of the model, which assumes the existence of three cellular types (*P* : progenitors, *D*: differentiated cells, *L*: differentiating cells committed to cell cycle completion). *P* cells differentiate with a probability *q*; otherwise they continue proliferating. Differentiating cells become either completely (*D*) with probability *δ* or commit to cell cycle completion (*L*) with probability 1 *− δ*. All *L* cells complete the cellular cycle and generate two *D* cells. **B)** The probability *q* of differentiation dictates whether the pool of progenitors expands and drives exponential tissue expansion (*q <* 1*/*2, left panel); remains steady and (*q* = 1*/*2, middle panel) and induces steady tissue growth at a constant pace; or vanishes and provokes tissue maturation, i.e., cease of growth (*q >* 1*/*2, right panel). These regimes are independent of the parameter *δ*. **C)** For expanding systems (*q <* 1*/*2), the model predicts that the *Q* fraction (the proportion of newly formed *D* cells) will remain constant over time (left panel) and this will produce a long-term accumulation of differentiated cells which will reach 2*q* in proportion (right panel).

- self-renewing progenitor cells (*P*), which constantly proliferate and differentiate;
- terminally differentiated cells (*D*), which irreversibly exit the cell cycle (i.e., become post-mitotic); and
- cells that have initiated the differentiation program but will undergo one final division round before exiting the cell cycle. We will call these late-differentiated cells *L* cells (see Fig. 2A).

Each newly formed cell from a dividing progenitor *P* will adopt one of the three fates above with certain probabilities: a cell might differentiate with probability *q* or remain as a *P* cell with probability 1 *− q*. Furthermore, a differentiating cell might either adopt *N* or *L* fate with probabilities *δ* and 1 *− δ*, respectively (Figure 2A). All *L* cells eventually divide generating two *N* cells. As we will see, it is this last premise the one responsible of breaking the observed statistical association between cell division and cell fate. This two-parameter model is general enough to encompass two extreme scenarios: i) independent fate assignment of daughter cells would correspond the case were *δ* = 1; and ii) development through purely symmetric divisions is attained when *δ* = 0. All intermediate situations are thus captured by values of *δ* between zero and one.

In a first approximation we disregard the variability in the cycle length of proliferating cells and thus only consider discrete generations of cells. In this scenario, the mean behaviour of the cell populations can be described by the following system of linear first order recurrence relations:

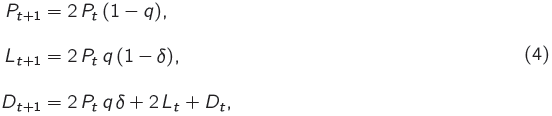

where *P*_*t*_, *L*_*t*_ and *D*_*t*_ are the average numbers of cells of type *P*, *L* and *D* in the *t*-th generation, respectively. *N*_*t*_ on the other hand accounts for the newly generated post-mitotic cells in the *t*-th generation, which, in the absence of cell loss, accumulate as *D* cells.

The above system of equations can be analytically solved. For *q >* 1*/*2 the number of progenitor cells *P*_*t*_ will decrease exponentially and the system will eventually deplete from both *P* and *L* cells. For *q* = 1*/*2 the system will be in a critical state, with a constant average number of progenitor cells and, in the absence of cell loss, the pool of differentiated cells will grow linearly. Finally, as expected, for *q <* 1*/*2 the system will growth exponentially. In this case the solution for the evolution of a monoclonal population for (*P*_0_ = 1*, L*_0_ = 0*, N*_0_ = 0*, D*_0_ = 0) is:

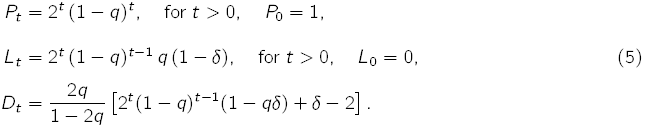

To that end we shall solve the long term proportions of the cell pools and find the ratios of each division mode. Two magnitudes of interest are the total number of newly formed cells in each generation, *F*_*t*_ = *P*_*t*_ + *L*_*t*_ + *N*_*t*_, and the total number of differentiated cells, *T*_*t*_ = *P*_*t*_ + *L*_*t*_ + *D*_*t*_ :

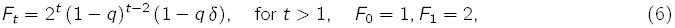

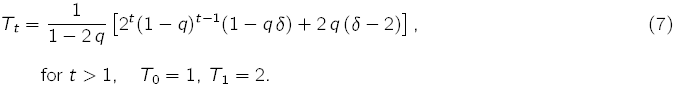

With these we can define the proportions of newborn cells in each generation as: For 0 *< q <* 1*/*2 the model predicts that the proportions of cell types in the newly formed cells are constant over time:

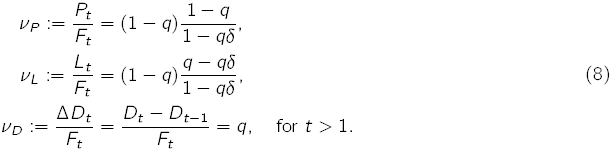

The fraction of newborn cells that differentiated into *D* cells (*?_D_*) is exactly *q* despite each cell stemming from a progenitor cell has a probability *qδ* of becoming a differentiated post-mitotic cell (Fig. 2A). In other words, the contribution of dividing *L* cells to the pool of post-mitotic cells compensates for the fact that not all differentiating cells become *D* cells. The number of generated cells at each generation is exactly twice the number of *P* and *L* cells in the previous generation, as expected:

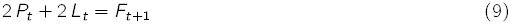

Thus, the model parameter *q* matches exactly the definition of the *Q* fraction stated above and all pools of cells will expand following a geometric progression with a growth rate that depends on the probability of differentiation *q*.

And a similar result can be obtained for the total proportions of cells in the long term (for steady growth
conditions 0 *<q <* 1*/*2):

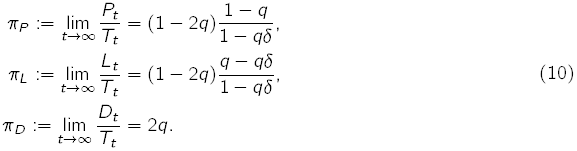

Both the proportion of newly generated *D* cells, *?_D_* = *q* and the total number of *D* cells accumulated, *π*_*D*_ = 2*q* are also defined only with the parameter *q*. It is worth noticing that the growth ratio for the total number of cells produced as well as in each pools of cells is 2(1 *− q*) and thus does not depend on the parameter *δ*. In fact, the model predicts that the number of cells will double every log(2)*/*(1 *− q*) cell cycles (see appendix).

With these proportions of we can already calculate the fraction of each division type. In order to appropriately define the three division modes we make the distinction between proliferating (*P* and *L*) and non-proliferating (*D*) cells and classify the seven division events of our model accordingly, as shown in Figure 3A. The Figure also indicates the probabilities of each division type for each division of a *P* or *L* cell.

**Figure 3.**
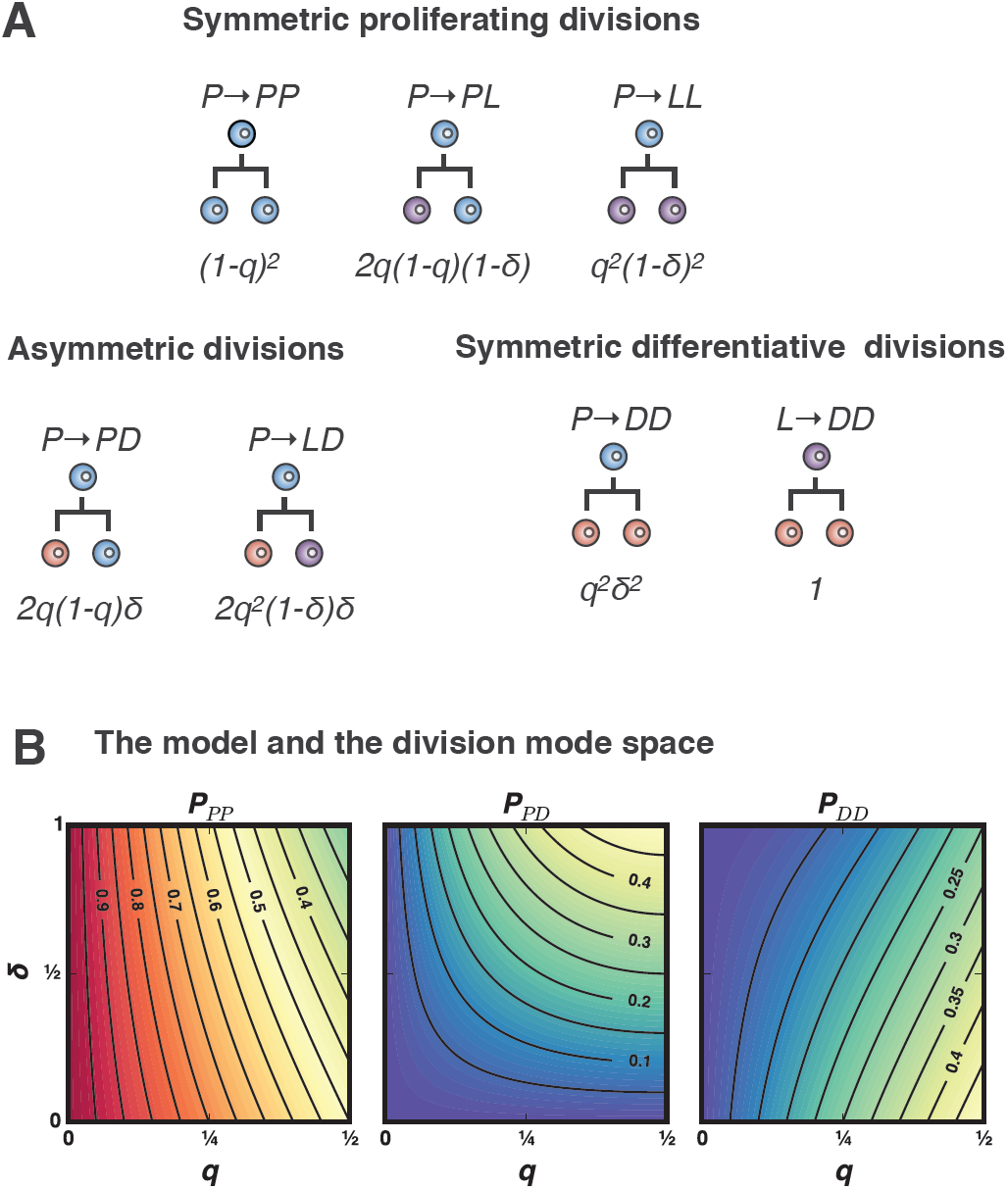
The model can recapitulate all division mode stragegies. **A)** The model entails 7 distinct division modes involving *P*, *L* and *D* cells. For the purpose of comparison with experimental data, these are grouped into symmetric, *P D* and *DD*, depending on whether the progeny is proliferative or post-mitotic. **B)** The model recapitulates all growth regimes (*q ≤*1*/*2) that involve an excess of symmetric divisions (0 *≤ δ ≤* 1, cf. middle panel with third quadrant of Fig. 1C).

With these and the proportions calculated above, the probabilities of each

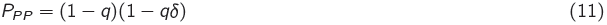

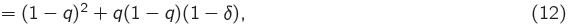

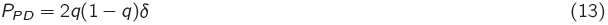

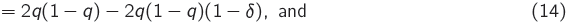

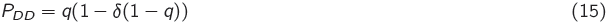

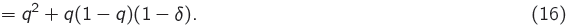

From these equations it is obvious that *δ* is just a shifted version of the divergence *δ* = *δ −* 1 (see Fig. 2C) and therefore our model spans the negative range of *δ*, in other words, the region of the division mode space where symmetric divisions are enriched (i.e., the third quadrant in Fig.1)

Another implication of (11) is that the two parameters of the model can be estimated from the experimental data:

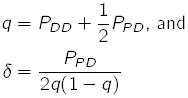

We have already seen that the rate of exponential expansion depends on the probability of differentiation (2(1 *− q*)) but not on the divergence. Nevertheless, the total number of differentiated cells is affected by the amount of symmetric divisions. Figure 2A exemplifies this by comparing two growth strategies with the same probability of differentiation (*q* = 0.4) and different proportions of symmetric divisions (*δ* = 0.1 vs. *δ* = 0.9). Clearly, the larger the number of symmetric divisions, the larger the total number of differentiated cells. According to (5), the fold-change increase in the number of *D* cells in the system with respect to the equilibrium case for a given *δ* is 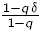(Fig. 4) Hence, abundance of symmetric divisions promotes more efficient generation of differentiated cells, allowing for up to a 2-fold increase in systems with high differentiation rates.

## Discussion

Here we have addressed the impact that different strategies of progenitor cell division have on the growth of organs and tissues. We provide an account of the possible modes of growth and maturation from symmetric and asymmetric cell divisions, and propose the use of two parameters: the probability of differentiation *q* (known as the *Q* fraction), and the divergence from the *linkage equilibrium* situation, *δ* (i.e., that in which daughter cells differentiate independently of each other) as general descriptors of growth and differentiation dynamics. We focus our attention on growing systems in which the number of symmetric cell divisions exceeds that expected by chance if the fate of sibling cells was uncorrelated, and propose a neutral model to account for the incidence of this excess. The model predicts that, given a fixed probability of differentiation, the divergence towards symmetric divisions has no impact in the exponential growth rate (i.e., in the cell number doubling time), but that it impacts the efficiency of generating differentiated cells. We observe that the higher the proportion of symmetric divisions, the more efficient is the deployment of differentiated cells.

**Figure 4.**
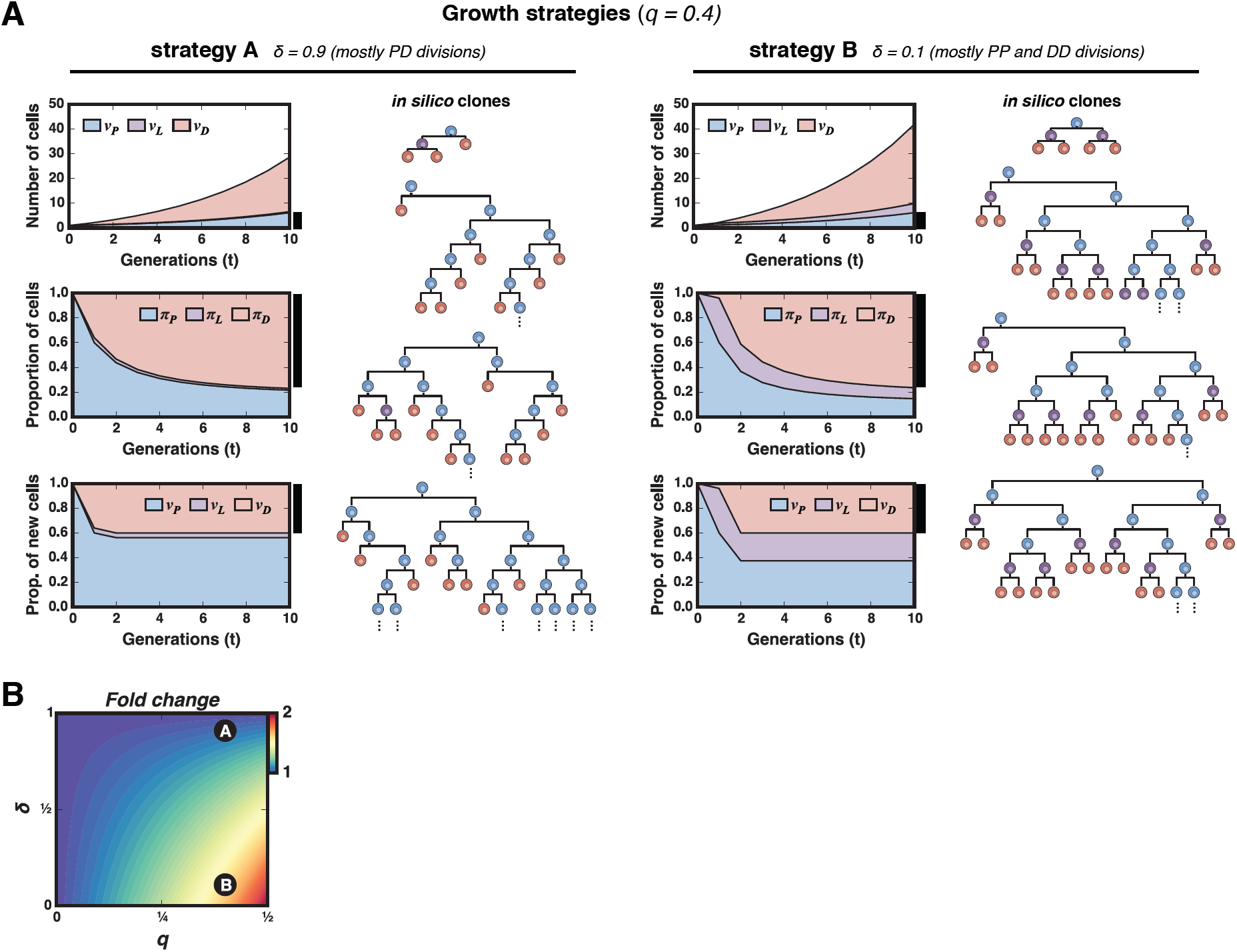
Symmetric divisions allow for more efficient generation of differentiated cells. A bias towards symmetric cell divisions (*δ <* 1) produces more efficient deployment of differentiated cells. **A)** Two strategies of growth with the same probability of differentiation are depicted. Strategy 1 (left panels, *δ* = 0.9) consists of progenitors dividing nearly in equilibrium, i.e., daughter cells differentiating almost completely independently of each other. Strategy 2 (right panels, *δ* = 0.9) comprises progenitors undergoing mainly *P P* and *DD* divisions, the differentiated cells in this case outnumber the *D* cells generated with strategy 1. Under identical differentiation conditions (same *q* and thus same growth rate), the most efficient growth strategy to maximize the yield in the differentiated pool consists in increasing the proportion of symmetric cell divisions. This is illustrated with the representative simulated clones displayed for each of the two strategies. With symmetric divisions (strategy 2) we almost double the yield of differentiated cells. **B)** The effect of increased symmetric divisions in the generation of differentiated cells is stronger the slower the system is growing. The colormap displayed shows the fold-change increase in differentiated cells with respect to the case where sibling cells differentiated independently.

There are many known examples of developmental systems using different and dynamic strategies of cell division (Gunage, Reichert, and VijayRaghavan 2014; Itzkovitz et al. 2012; Livesey and Cepko 2001). In all cases it has been suggested that the optimal growth program, in terms of minimum developing time, requires a first phase of exponential proliferation of stem cells through symmetric divisions, followed by a phase of asymmetric divisions that generate the differentiated cells (Itzkovitz et al. 2012). This strategy has been successfully tested in the context of the development of mouse intestinal crypts (Itzkovitz et al. 2012) and also matches quite well the observed behaviour of progenitors in during the development of the CNS (Livesey and Cepko 2001). The fact that adult organ sizes are constant within animal species indicates that these growth master plans implement robust size specification by means of tight regulation of proliferation and differentiation, which might involve feedback controls, as it has also been suggested (Lander et al. 2009). These studies on tissue and organ growth mainly focus on the regulation of the rate or probability of differentiation, but not on the specific strategies of cell divisions (i.e., the divergence) and the impact that these may have on the developing system. We believe that positioning these questions in the context of the divergence parameter proposed in this work together with the estimates of differentiation probabilities will provide insights into how cell division is related to the controlled growth of a developing tissue.

On other hand, the impact of the division modes on the dynamics of individual clones has been extensively
studied in the context of homeostatic systems (Clayton et al. 2007; Klein, Nakagawa, et al. 2010; Simons and Clevers 2011; Snippert et al. 2010), where the probability of differentiation is roughly 50%. Clonal analysis in these systems leads to a model of stochastic cell turnover sustained by the random loss and replacement of stem cells (Klein and Simons 2011). In this model, the cell population is maintained steadily by a tight balance between loss and proliferation (probability of differentiation is assumed to be constant and equal to 1/2). According to the theory, the stochastic nature of divisions should imprint the clone statistics with a signature of a neutral competition; namely, the constant loss of clones due to accidental extinction of progenitors, the ever-broadening distribution of clone sizes, or the takeover of some clones over others by chance (Klein and Simons 2011). Recently, an extrapolation of these models to developmental systems has been used to account for the emergence of the retina in zebrafish (He et al. 2012). The proposed model, based on stochastic cell division and fate assignation reproduces the evolution of the distribution of clonal sizes over time. However, this model, which does not take into account the well established sequence of events associated with the development of this organ, overlooks the changes in the division modes and its association to changes in the cell cycle. In fact, during embryonic neurogenesis the transition from proliferation to differentiation is concomitant with the lengthening of the cell cycle (Calegari and Huttner 2003). This has led to the *cell cycle length hypothesis* (Calegari and Huttner 2003; Lange, Huttner, and Calegari 2009) that states that the length of G1 phase impacts differentiation by permitting fate determinants to achieve its neurogenic effect (Götz and Huttner 2005).

Other developmental systems show a similar progression pattern to the nervous system: an early exponential amplification phase of the progenitor pool by means of symmetric division followed by a period of asymmetric divisions and a final phase of terminal symmetric divisions (Egger, Gold, and Brand 2010; Gunage, Reichert, and VijayRaghavan 2014). This behaviour can be qualitatively accounted by an increasing rate of differentiation *q*. However, the particular biases in the division modes can only be accounted for through specific mechanisms. Here we propose a framework for its analysis and a basic –or neutral– model that accounts for such biases.

Such hypothesis could be posed in the terms proposed in our framework. The temporal changes in the division modes can be captured not only by the *Q* fraction but also by the divergence *δ*. One possible interpretation for the changes in *δ* induced by changes in the cell cycle length is depicted in Figure 5: as undifferentiated cells progress through the cell cycle these might decide to terminally differentiate at varying times. We shall call this the differentiation event. If this event occurs soon after division (red section in Fig. 5), where the new cell has started its progression through G1, it is conceivable that this cell might exit the cycle and become post-mitotic. On the other hand, if the differentiation event occurs later on during the cell cycle (purple section in the Fig. 5), the cell might have already committed to the completion of the cellular cycle and thus shall undergo a last division round that will give rise to two differentiated cells. In this scenario, cells committing to differentiation once they are already committed to cell cycle completion, will contribute extra symmetric terminal divisions. The relative proportion between the time undifferentiated cells spend in the early (red) and late (purple) phases will dictate the magnitude of the divergence and the excess of symmetric divisions in the system. Therefore, a change in the cell cycle length could lead to a change in the division mode even in the case where the probability of differentiation *q* remains unaltered. Of course, observations indicate that the differentiation rate increases as the cell cycle lengthens and this directly impacts the division mode. Our model can thus be used to distil the contribution of both effects, the increase in the differentiation rate per se and the lengthening of the cycle).

**Figure 5.**
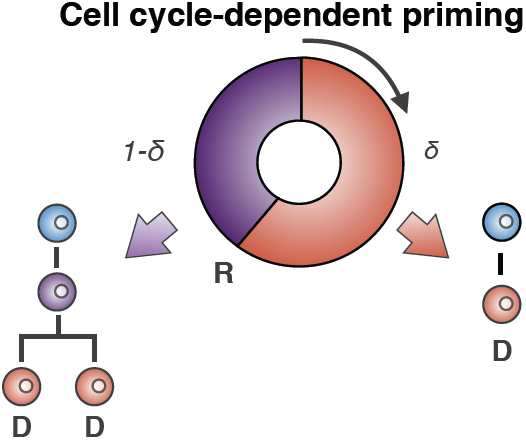
Cell cycle-dependent commitment to differentiation could explain the distribution of symmetric vs. asymmetric divisions. The observed abundance of symmetric differentiative divisions could be explained by the interplay between the cell cycle and differentiation. Cells committing to differentiation after having passed a point of no return (R) in the cell cycle shall divide symmetrical into two differentiated cells.

Together, the framework presented here for the analysis of expansion and differentiation in developing systems and the proposed neutral model provides an initial step towards a framework for the analysis of specific mechanisms driving the changes in the proliferative behaviour of progenitor cells.

## References

Blanpain, C. and B. D. Simons (2013). “Unravelling stem cell dynamics by lineage tracing.” In: Nature reviews. Molecular cell biology 14.8, pp. 489–502. DOI: 10.1038/nrm3625.

Bultje, R. S., D. R. Castaneda-Castellanos, L. Y. Jan, Y.-N. Jan, A. R. Kriegstein, and S.-H. Shi (2009). “Mammalian Par3 regulates progenitor cell asymmetric division via notch signaling in the developing neocortex.” English. In: Neuron 63.2, pp. 189–202. DOI: 10.1016/j.neuron.2009.07.004.

Calegari, F. and W. B. Huttner (2003). “An inhibition of cyclin-dependent kinases that lengthens, but does not arrest, neuroepithelial cell cycle induces premature neurogenesis”. In: Journal of Cell Science 116.24, pp. 4947–4955. DOI: 10.1242/jcs.00825.

Caviness, V., T. Takahashi, and R. Nowakowski (1995). “Numbers, time and neocortical neuronogenesis: a general developmental and evolutionary model”. In: Trends in Neurosciences 18.9, pp. 379–383. DOI: 10.1016/0166-2236(95)93933-0.

Centanin, L., J.-J. Ander, B. Hoeckendorf, K. Lust, T. Kellner, I. Kraemer, C. Urbany, E. Hasel, W. A. Harris, B. D. Simons, and J. Wittbrodt (2014). “Exclusive multipotency and preferential asymmetric divisions in post-embryonic neural stem cells of the fish retina.” In: Development (Cambridge, England) 141.18, pp. 3472–3482. DOI: 10.1242/dev.109892.

Clayton, E., D. P. Doupé, A. M. Klein, D. J. Winton, B. D. Simons, and P. H. Jones (2007). “A single type of progenitor cell maintains normal epidermis.” In: Nature 446.7132, pp. 185–9. DOI: 10.1038/ nature05574.

Egger, B., K. S. Gold, and A. H. Brand (2010). “Notch regulates the switch from symmetric to asymmetric neural stem cell division in the Drosophila optic lobe.” In: Development (Cambridge, England) 137.18, pp. 2981–7. DOI: 10.1242/dev.051250.

Fisher, R. A. (1922). “On the Interpretation of X2 from Contingency Tables, and the Calculation of P.” en. In: Journal of the Royal Statistical Society 85.1, pp. 87–94.

Gönczy, P. (2008). “Mechanisms of asymmetric cell division: flies and worms pave the way.” In: Nature reviews. Molecular cell biology 9.5, pp. 355–66. DOI: 10.1038/nrm2388.

Götz, M. and W. B. Huttner (2005). “The cell biology of neurogenesis.” In: Nature reviews. Molecular cell biology 6.10, pp. 777–88. DOI: 10.1038/nrm1739.

Gunage, R. D., H. Reichert, and K. VijayRaghavan (2014). “Identification of a new stem cell population that generates Drosophila flight muscles”. In: eLife 3. Ed. by D. Pan.

Hara, K., T. Nakagawa, H. Enomoto, M. Suzuki, M. Yamamoto, B. D. Simons, and S. Yoshida (2014). “Mouse spermatogenic stem cells continually interconvert between equipotent singly isolated and syncytial states.” English. In: Cell stem cell 14.5, pp. 658–72. DOI: 10.1016/j.stem.2014.01.019.

Hardwick, L. J. A. and A. Philpott (2014). “Nervous decision-making: to divide or differentiate.” In: Trends in genetics : TIG 30.6, pp. 254–61. DOI: 10.1016/j.tig.2014.04.001.

Hardy, G. H. (1908). “Mendelian Proportions in a Mixed Population”. In: Science.

He, J., G. Zhang, A. D. Almeida, M. Cayouette, B. D. Simons, and W. A. Harris (2012). “How variable clones build an invariant retina.” In: Neuron 75.5, pp. 786–98. DOI: 10.1016/j.neuron.2012.06.033.

Itzkovitz, S., I. C. Blat, T. Jacks, H. Clevers, and A. van Oudenaarden (2012). “Optimality in the development of intestinal crypts.” In: Cell 148.3, pp. 608–19. DOI: 10.1016/j.cell.2011.12.025.

Kim, Y. H., H. L. Larsen, P. Rué, L. A. Lemaire, J. Ferrer, and A. Grapin-Botton (2014). “Cell cycle-dependent differentiation dynamics balances growth and endocrine differentiation in pancreas”. In: (in preparation).

Klein, A. M., T. Nakagawa, R. Ichikawa, S. Yoshida, and B. D. Simons (2010). “Mouse germ line stem cells undergo rapid and stochastic turnover.” In: Cell stem cell 7.2, pp. 214–24. DOI: 10.1016/j.stem.2010. 05.017.

Klein, A. M. and B. D. Simons (2011). “Universal patterns of stem cell fate in cycling adult tissues.” In: Development (Cambridge, England) 138.15, pp. 3103–11. DOI: 10.1242/dev.060103.

Knoblich, J. A. (2008). “Mechanisms of asymmetric stem cell division.” In: Cell 132.4, pp. 583–97. DOI: 10.1016/j.cell.2008.02.007.

Kretzschmar, K. and F. M. Watt (2012). “Lineage tracing.” In: Cell 148.1-2, pp. 33–45. DOI: 10.1016/j. cell.2012.01.002.

Lander, A. D., K. K. Gokoffski, F. Y. M. Wan, Q. Nie, and A. L. Calof (2009). “Cell lineages and the logic of proliferative control.” In: PLoS biology 7.1. Ed. by C. F. Stevens, e15. DOI: 10.1371/journal.pbio. 1000015.

Lange, C., W. B. Huttner, and F. Calegari (2009). “Cdk4/cyclinD1 overexpression in neural stem cells shortens G1, delays neurogenesis, and promotes the generation and expansion of basal progenitors.” In: Cell stem cell 5.3, pp. 320–31. DOI: 10.1016/j.stem.2009.05.026.

Lescroart, F. S. Chabab, X. Lin, S. Rulands, C. Paulissen, A. Rodolosse, H. Auer, Y. Achouri, C. Dubois, A. Bondue, B. D. Simons, and C. Blanpain (2014). “Early lineage restriction in temporally distinct populations of Mesp1 progenitors during mammalian heart development.” In: Nature cell biology 16.9, pp. 829–840. DOI: 10.1038/ncb3024.

Li, H.-S., D. Wang, Q. Shen, M. D. Schonemann, J. A. Gorski, K. R. Jones, S. Temple, L. Y. Jan, and Y. N. Jan (2003). “Inactivation of Numb and Numblike in Embryonic Dorsal Forebrain Impairs Neurogenesis and Disrupts Cortical Morphogenesis”. In: Neuron 40.6, pp. 1105–1118. DOI: 10.1016/S0896-6273(03) 00755-4.

Livesey, F. J. and C. L. Cepko (2001). “Vertebrate neural cell-fate determination: lessons from the retina.” In: Nature reviews. Neuroscience 2.2, pp. 109–18. DOI: 10.1038/35053522.

McConnell, S. K. (1995). “Constructing the cerebral cortex: Neurogenesis and fate determination”. In: Neuron 15.4, pp. 761–768. DOI: 10.1016/0896-6273(95)90168-X.

Noctor, S. C., V. Martínez-Cerdenõ, L. Ivic, and A. R. Kriegstein (2004). “Cortical neurons arise in symmetric and asymmetric division zones and migrate through specific phases.” In: Nature neuroscience 7.2, pp. 136–44. DOI: 10.1038/nn1172.

Nomura, T., H. Gotoh, and K. Ono (2013). “Changes in the regulation of cortical neurogenesis contribute to encephalization during amniote brain evolution.” en. In: Nature communications 4, p. 2206. DOI: 10.1038/ncomms3206.

Rakic, P. (1995). “A small step for the cell, a giant leap for mankind: a hypothesis of neocortical expansion during evolution”. In: Trends in Neurosciences 18.9, pp. 383–388. DOI: 10.1016/0166-2236(95)93934P.

Rapaport, D. H., L. L. Wong, E. D. Wood, D. Yasumura, and M. M. LaVail (2004). “Timing and topography of cell genesis in the rat retina.” In: The Journal of comparative neurology 474.2, pp. 304–24. DOI: 10.1002/cne.20134.

Simons, B. D. and H. Clevers (2011). “Strategies for homeostatic stem cell self-renewal in adult tissues.” English. In: Cell 145.6, pp. 851–62. DOI: 10.1016/j.cell.2011.05.033.

Snippert, H. J., L. G. van der Flier, T. Sato, J. H. van Es, M. van den Born, C. Kroon-Veenboer, N. Barker, A. M. Klein, J. van Rheenen, B. D. Simons, and H. Clevers (2010). “Intestinal crypt homeostasis results from neutral competition between symmetrically dividing Lgr5 stem cells.” In: Cell 143.1, pp. 134–44. DOI: 10.1016/j.cell.2010.09.016.

Takahashi, T., R. S. Nowakowski, and J. Caviness Verne S. (1996). “The Leaving or Q Fraction of the Murine Cerebral Proliferative Epithelium: A General Model of Neocortical Neuronogenesis”. In: J. Neurosci. 16.19, pp. 6183–6196.

